# Snake Kolmiovirus Encodes a Single Form of Delta Antigen and Shows No Evidence of Translation from Open Reading Frame 2

**DOI:** 10.64898/2026.04.10.717655

**Authors:** Leonóra Szirovicza, Udo Hetzel, Tomas Strandin, Anja Kipar, Jussi Hepojoki

**Author notes:** Corresponding author: Correspondence should be addressed to Leonóra Szirovicza, *Mailing address*: University of Helsinki, Department of Virology, P.O. Box 21, Haartmaninkatu 3, FI-00014 University of Helsinki, Finland.

## Abstract

Hepatitis D virus (HDV) is a satellite virus that utilizes hepatitis B virus (HBV) as a helper for infectious particle formation. HDV was originally identified as a novel antigen in liver biopsies of HBV patients, and later studies showed the “delta” antigen (DAg) to be the sole protein encoded by HDV. Until the discovery of HDV-like agents in birds and snakes in 2018, HDV was a unique example of animal satellite viruses. We identified Swiss snake colony virus 1 (SwSCV-1) in the brain of a *Boa constrictor*, and through comparison we found the genome organization of SwSCV-1 to resemble that of HDV. However, in addition to the DAg open reading frame (ORF), the genome of SwSCV-1 includes another >500 nt ORF, “ORF2”. To study whether the putative ORF2-encoded protein plays a role in the SwSCV-1 life cycle, we established an infectious clone of the virus with a point mutation in the methionine initiation codon of ORF2. The mutation did not significantly affect initiation of replication, establishment of persistent infection, or infectious particle formation upon superinfection with a helper virus. Using additional methods, we gathered further evidence confirming that ORF2 is not actively translated in boa constrictor cells. We further showed that unlike HDV, SwSCV-1 expresses a single form of the DAg. Although the proteins encoded by SwSCV-1 and HDV only include one and two forms of the DAg, respectively, whether other kolmioviruses express additional forms of DAg or related proteins in some cell types or host species merits further research.

**IMPORTANCE:** Approximately 40 years after the discovery of hepatitis D virus (HDV), satellite viruses with similar genome organization were found in various animals, thereby giving rise to family *Kolmioviridae*. HDV encodes a single protein, the delta antigen (DAg), which comes in small and approximately 20 amino acids longer large form. The genome of some HDV species and many of the newly found kolmioviruses contains additional open reading frames (ORFs), potentially enabling protein expression. Here, we studied the viral proteins expressed during Swiss snake colony virus 1 (SwSCV-1) infection of boa constrictor cells. Our findings show that unlike HDV, SwSCV-1 encodes only a single form of DAg. In addition, our study suggests that, like in HDV, the additional ORF in SwSCV-1 genome does not give rise to a protein. Although we could not demonstrate expression of additional viral proteins during SwSCV-1 infection, it is important to study the proteome of other kolmioviruses.

## INTRODUCTION

Hepatitis D virus (HDV) – first described in 1977 – is a unique pathogen, originally identified in liver biopsies of hepatitis B virus (HBV) carriers (1). Soon after its discovery, chimpanzee experiments allowed the researchers to recognize HDV as a satellite virus of HBV, needing the glycoproteins of the latter for infectious particle formation (2, 3). HDV has a single-stranded negative-sense circular RNA genome of ∼1.7 kilobases, which – similarly to viroids – replicates via a double rolling circle mechanism (4, 5). Three different HDV RNA species are present in the cell during its replication cycle: the genome and the antigenome, which are exact complements of each other, and the ∼800 nucleotides long mRNA, which gives rise to the delta antigen (DAg) (5). During HDV replication, multimeric RNA species are generated; these are processed into unit-length monomers by ribozymes that are present in both the genomic and antigenomic RNAs (6). In case of HDV, the DAg exists in two forms (small [S-DAg] and large [L-DAg]), due to the RNA editing activity of the cellular enzyme adenosine deaminase acting on RNA (ADAR) (7–9). As the consequence of the ADAR editing on the antigenomic HDV RNA strand (10), the stop codon of the DAg ORF is modified into a tryptophan codon, resulting in elongation of the DAg by 19 additional amino acids to generate the L-DAg (11). The S-DAg promotes replication, while the L-DAg has an inhibitory effect on the replication and shifts the cycle towards particle formation (12). Post-translational modifications such as phosphorylation, acetylation, and isoprenylation likely contribute to the vastly different roles that the two forms of the DAg play during HDV replication (12, 13).

For more than 40 years, HDV was the sole representative of the unassigned genus *Deltavirus* (14). In 2018, coincidentally and independently to Wille and colleagues, who identified a “novel” HDV-like agent in ducks (15), we identified an HDV-like agent in the brain of a *Boa constrictor* (16). In both cases, the HDV-like virus appeared to exist without a co-infecting hepadnavirus, and in fact we later demonstrated reptarena- and hartmaniviruses to serve as the helper viruses for the “snake deltavirus”, i.e. Swiss snake colony virus 1 (SwSCV-1) (17). Subsequent reports from different groups described additional HDV-like agents in termites, fish, frogs, newts, rodents, bats, deer, woodchucks and birds (18–21), which provoked the International Committee on Taxonomy of Viruses to establish the realm *Ribozyviria* and the family *Kolmioviridae* that currently includes eleven genera (22–24).

While the characterization of the newly identified kolmiovirids is in its infancy, it appears likely that these agents share many of HDV’s characteristics. The HDV genome contains several ORFs, however, the evidence suggests that the two forms of the DAg are the only proteins translated (5, 25). In addition to the DAg ORF, the genome of SwSCV-1 contains a second ORF (ORF2) in the genomic orientation that would give rise to a putative protein of 177 amino acids (16). While not studied for most of the novel kolmioviruses (15, 18–21), a recent study found transcriptomics-based evidence of the presence of mRNA from a genomic-orientation ORF in the *Serinus canaria*-associated deltavirus genome (26). The researchers did not identify significant homologies to any known protein for the *Serinus canaria*-associated deltavirus “ORF2” product (26), similar to the SwSCV-1 ORF2 protein product (16).

In this study, we investigated whether the SwSCV-1 ORF2 and its putative protein product play a role in the life cycle of the virus. We first introduced a mutation to remove the ORF2 methionine initiation codon while retaining the intact DAg ORF in the plasmid-driven SwSCV-1 infectious clone. We then compared replication initiation, infectious particle formation and persistence of infection between the ORF2-deleted and wild-type (2× and 1.2× SwSCV-1) genome clones in snake cells. Next, we utilized recombinant protein expression to study the cellular localization of the putative ORF2 encoded protein, and to demonstrate that only a single form of the DAg is produced in SwSCV-1 infected boid cells. Finally, to rule in or out the presence of ORF2 mRNA, we performed directional RNA sequencing utilizing two approaches (rRNA depletion and poly(A) enrichment) for library preparation from RNA extracted from SwSCV-1 infected cells.

## MATERIALS AND METHODS

### Plasmids and cloning

The pCAGGS plasmid-based infectious clones bearing either 2× or 1.2× SwSCV-1 genome are described in (17, 27). We ordered synthetic genes with 1.2× genome copies of SwSCV-1 (accession number: NC_040729.1), with the methionine initiation codon of the ORF2 mutated to isoleucine (ATG→ATA) leaving the DAg ORF unaffected (**Figure 1A**). In addition, we ordered synthetic genes representing the putative protein translated from ORF2 with and without a stop codon. The absence of the stop codon allows attachment of an HA-tag (encoded in the vector) to the translated protein (**Figure 1B**) and subsequent detection with the tag specific antibody. In addition, we ordered three synthetic genes for the DAg ORF; one with the ORF extending only to the S-DAg stop codon, one with the natural ORF until the end of the putative L-DAg stop codon, and one with the genome-encoded stop codon altered to tryptophan codon (TAG→TGG) to force L-DAg expression (**Figure 1C**). All synthetic genes were made by GeneUniversal. We subcloned the inserts into pCAGGS-MCS (28) following the protocol described in (17). In brief, after FastDigest (ThermoFisher Scientific) restriction digestion with suitable enzymes and agarose gel electrophoresis of the inserts, the GeneJET Gel extraction kit (ThermoFisher Scientific) served to purify the products. After linearization, and blunting in the case of the infectious clones, of the pCAGGS/MCS (28) plasmid, T4 DNA ligase (ThermoFisher Scientific) served to ligate the plasmid and the inserts. We used the ligation products to transform chemically competent *E. coli* (DH5α strain) and subsequently plated the cells on Luria-Broth (LB) agar plates with 100 µg/ml of ampicillin, and incubated overnight (O/N) at 37 °C. We picked and transferred single colonies into 5 ml of LB medium (10 g/l tryptone, 10 g/l NaCl, 5 g/l yeast extract), incubated them O/N at 37 °C at 220 rpm, and isolated plasmids using the GeneJET Plasmid Miniprep Kit (ThermoFisher Scientific) from 2 ml of the O/N culture. Sanger sequencing (provided by the DNA Sequencing and Genomic Laboratory, Institute of Biotechnology, University of Helsinki) confirmed the presence of the correct inserts. ZymoPURE II Plasmid Maxiprep Kit (Zymo Research) following the manufacturer’s protocol served for preparing endotoxin-free plasmid stocks.

**Figure 1.**
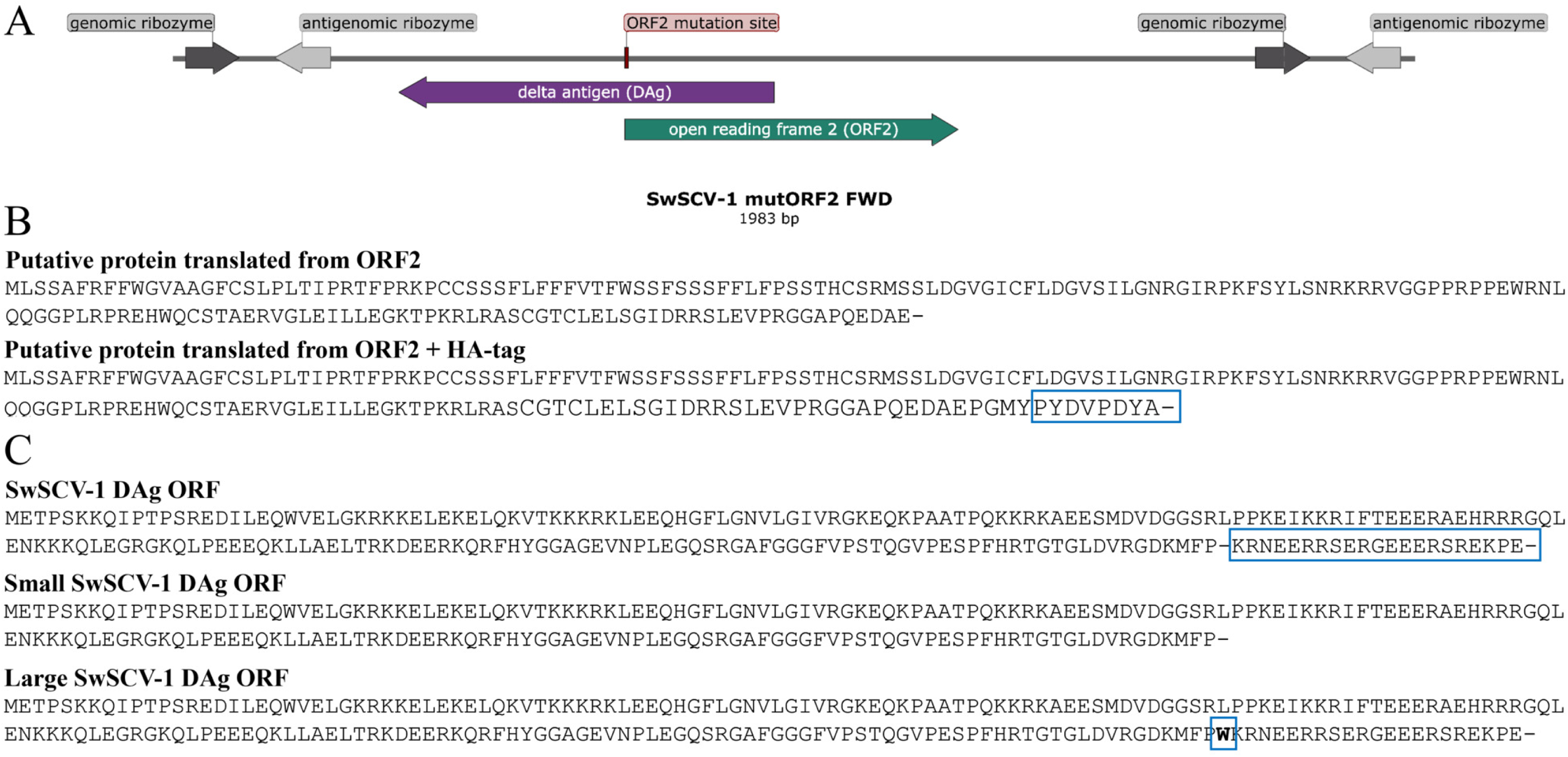
Schematic representation of the synthetic genes and protein products. **A)** Schematic representation of the 1.2× genome SwSCV-1 insert in forward (FWD)/genomic orientation with the introduced point mutation in the open reading frame 2 (ORF2). The point mutation (ATG→ATA) in the ORF2 initiation codon inhibits translation of the putative protein, while leaves the delta antigen unaffected. **B)** Amino acid sequence of the ORF2 protein translated with or without an HA-tag (marked with a blue rectangle) from the expression construct. **C)** Amino acid sequence of the proteins translated from the synthetic genes representing the three forms of the SwSCV-1 delta antigen (DAg): the full unedited version, allowing RNA editing (site marked by the blue rectangle); the small form with no possibility for editing and extension; and the edited DAg (editing site marked by the blue square), forcing the expression of the large form of the protein.

### Cell culture and superinfection

The study made use of a *Boa constrictor* kidney cell line, I/1Ki, spontaneously immortalized from kidney homogenate (29); and two persistently SwSCV-1 infected cell lines, I/1Ki-Δ and I/1Ki-1.2×Δ (17, 27). To generate the I/1Ki-mutORF2Δ cell line, we transfected I/1Ki cells with the 1.2× SwSCV-1 genome clone with disabled ORF2, followed by continuous passaging and maintenance of the cells. In addition to the abovementioned cell lines, we utilized four persistently SwSCV-1 infected cell lines (V/5Lu-Δ, V/1Ki-Δ, V/2Hz-Δ, and V/5Liv-Δ) described in (17). One further persistently SwSCV-1 infected cell line was established, designated V/4Br-Δ. Briefly, we transfected previously described (30) boa constrictor brain cells (V/4Br) with the 1.2× SwSCV-1 FWD infectious clone (27), followed by continuous passaging of the cells as described for the above mentioned Δ cell lines (17).

In addition to the snake cell lines, we used human hepatocellular carcinoma (Huh-7) and human embryonic kidney (HEK293T) cells. Both cell lines were obtained from ATCC.

We maintained all snake cell lines in Minimal Essential Medium Eagle (Sigma Aldrich), Huh7 cells in Dulbecco’s modified Eagle’s medium (low glucose) (Sigma Aldrich) and HEK293T cells in Dulbecco’s modified Eagle’s medium (high glucose) (Sigma Aldrich). All medias were supplemented with 10% fetal bovine serum (Gibco), 200 mM L-glutamine (Sigma Aldrich), 100 µg/ml of streptomycin (Sigma Aldrich), and 100 U/ml of penicillin (Sigma Aldrich) in an incubator at 30 °C with 5% CO_2_.

For infectious particle formation studies of the persistently SwSCV-1 infected cell lines, we inoculated the cells with Haartman Institute snake virus (HISV-1) (31), shown to efficiently function as the helper virus of SwSCV-1 (17, 27). We used 600 copies of HISV-1 S segment RNA per cell as the infectious dose, which corresponds approximately to MOI (multiplicity of infection) 3 (32).

### Transfection

Lipofectamine 2000 (ThermoFisher Scientific) served as the reagent to transfect I/1Ki cells, as described (17, 33, 34). For Huh7 and HEK293T cells, the Fugene HD (Promega) reagent served for transfection purposes, as described in (34). Briefly, we prepared mixtures of 500 ng plasmid DNA in 50 µl OptiMEM (ThermoFisher Scientific), and 3 µl of Lipofectamine 2000 in 47 µl of OptiMEM (ThermoFisher Scientific). Then, we combined the mixtures by pipetting up and down and allowed complex formation for 15-30 min at room temperature (RT). Fugene HD was added directly into the DNA-OptiMEM mixture before allowing complex formation. We mixed 1 ml trypsinized cell suspension (∼ 1.8 cm^2^ of cells per ml, i.e. a confluent 75-cm^2^ suspended into 38 ml of fully supplemented medium) into the transfection reagent-plasmid mixture and allowed the suspension to stand at RT for 15-30 min before plating, and at 5-6 h post plating replaced the transfection mixture by fully supplemented growth medium. Depending on the upcoming experiment, we scaled up the above reaction volumes accordingly.

### Western blot (WB)

For WB, we scraped the cells into PBS, pelleted them by centrifugation (500 × g, 5 min), resuspended the pellets in RIPA buffer (50 mM Tris, 150 mM NaCl, 1% Triton-x-100, 0.1% sodium dodecyl sulfate [SDS], 0.5% sodium deoxycholate, protease inhibitor cocktail [Roche]), and measured the protein concentration of the samples using the Pierce^TM^ BCA Protein Assay Kit (ThermoFisher Scientific). We separated an equal amount of protein for each sample on SDS-PAGE using 4–20% Mini-PROTEAN® TGX gels (Bio-Rad), transferred onto nitrocellulose, and immunoblotted as described (35). We used rabbit α-SwSCV-1 DAg antiserum (16) as the primary antibody at 1:2000 dilution. For detection of proteins containing an HA-tag, a mouse anti-HA (clone 16B12; BioLegend) primary antibody was used at 1:1000 dilution. In addition, the serum of the boa constrictor, from which we originally identified SwSCV-1 (16), was also utilized for staining at 1:250 dilution, visualized by anti-IgY boa constrictor antibody (35) labelled using Alexa Fluor 680 NHS ester (ThermoFisher Scientic) following the manufacturer’s protocol. The other secondary antibodies used were the following: IRDye 680RD donkey anti-mouse (IgG) (LI-COR Biosciences) and IRDye 800CW donkey anti-rabbit (IgG) (LI-COR Biosciences), both at 1:10,000 dilution. Lab Vision^TM^ pan-actin mouse monoclonal antibody (ThermoFisher Scientific) at 1:200 dilution served to detect β-actin as a loading control, and the Odyssey Infrared Imaging System (LI-COR Biosciences) for recording the results.

### Immunofluorescence (IF) staining

For IF staining of cells, we followed a described protocol (17). Briefly, we plated the cells on collagen-coated CellCarrier-96 Ultra plates (PerkinElmer), fixed them with 4% paraformaldehyde (PFA) in PBS for ∼15 min at RT, incubated 15 min at RT in permeabilization buffer (50 mM Tris, 150 mM NaCl, 0.25% Triton-X-100, pH 7.4) supplemented with 3% bovine serum albumin to reduce unspecific biding and to quench the remaining PFA, and directly proceeded with staining. We used the directly labeled anti-SDAg-AF488 antibody described in (17) at 1:500 dilution or rabbit anti-SwSCV-1 DAg antiserum at 1:1,000 dilution. Alexa Fluor 488- or 647-labeled donkey anti-rabbit or anti-mouse immunoglobulin (ThermoFisher Scientific) at 1:1000 dilution served as the secondary antibodies. For the detection of proteins with an HA-tag, we used the mouse anti-HA (clone 16B12) primary antibody at 1:400 dilution. For imaging of the plates stored in the dark at 4 °C, we used the Opera Phenix High Content Screening System (PerkinElmer), provided by FIMM (Institute for Molecular Medicine Finland) High Content Imaging and Analysis (FIMM-HCA).

### Quantitative reverse transcription PCR (qRT-PCR)

qRT-PCR, described in (27), served for quantification of SwSCV-1 RNA in cells. RNAs isolated using the GeneJET RNA Purification Kit (ThermoFisher Scientific) served as the template for 10 µl reactions set up following the TaqMan® Fast Virus 1-Step Master Mix (ThermoFisher Scientific) protocol. The primers and probes used target the genomic strand of SwSCV-1. We ran the samples in duplicate using the AriaMX real-time PCR system (Agilent) with the following cycling conditions: reverse transcription for 5 min at 50 °C; initial denaturation for 20 s at 95 °C; and two amplification steps at 95 °C for 3 s and 60 °C for 30 s; repeated for 40 cycles. The TranscriptAid T7 High Yield Transcription Kit (ThermoFisher Scientific) served to generate *in vitro* transcribed SwSCV-1 specific control RNA which allowed us to generate a standard curve and to estimate the number of RNA copies in the samples (27). In addition, before assembling the qRT-PCR reactions, we subjected the samples to DNase I treatment (ThermoFisher Scientific) according to the manufacturer’s recommendations to eliminate any transfection leftover plasmid DNA in the RNA preparations.

We normalized the SwSCV-1 RNA levels against a house-keeping gene, using the following primers and probe for the detection of *Boa constrictor* phosphoglycerate kinase 1 (PGK-1): forward primer: 5’ GATGGATAAAGTTGTGGAA 3’, reverse primer: 5’ TTCGGTATTCCATTTAGC 3’, and probe 5’ 6-Fam (carboxyfluorescein)- TAGCAGTATCTCCGCCACCAAT -BHQ-1 3’. We used *Python bivitattus* PGK-1 (GenBank accession: XM_007429612.3) as the template in Unipro UGENE (36) for reference assembly to obtain *B. constrictor* PGK-1 mRNA from reads of our earlier metatranscriptomic studies (31, 37–39).

### Near-infrared fluorescent northern blot

Near-infrared northern blot analysis, performed as described in (27), served to detect viral RNA in the permanently SwSCV-1 infected cells. Briefly, we isolated RNA from I/1Ki, I/1Ki-2×Δ, I/1Ki-1.2×Δ and I/1Ki-mutORF2Δ cells by TRIzol^TM^ reagent (ThermoFisher Scientific), and separated the RNA samples in parallel with ssRNA ladder (New England Biolabs) on agarose-formaldehyde gels in tricine/triethanolamine buffer, as described by Mansour and Pestov (40). Following, we transferred the RNAs from the gel to Hybond^TM^-N^+^ nylon membrane (GE Healthcare) using O/N capillary transfer, then cross-linked the RNA to the membrane by an ultraviolet cross-linker (120 mJ/cm^2^ at 254 nm) (Analytik Jena). After, we prehybridized the membranes to block non-specific binding sites and performed O/N incubation with the fluorescent probes that were diluted into the pre-hybridization buffer to a final concentration of 1 nM. We used probes targeting the ladder, the SwSCV-1 genome and the DAg mRNA as described (27). The next day, we washed the membranes three times before recording the results utilizing the Odyssey Infrared Imaging System (LI-COR Biosciences).

### Next generation sequencing (NGS), strand specific RNAseq and Sanger sequencing of SwSCV-1 from infected cells

To investigate the SwSCV-1 genome within the infected cell, we performed metatranscriptomic analysis of the persistently SwSCV-1 infected cells. The NEBNext rRNA depletion kit (New England BioLabs) served for removal of ribosomal RNA from the total RNA isolated from cell pellets using the TRIzol^TM^ Reagent (ThermoFisher Scientific) according to the manufacturer’s recommendation. The NEBNext Ultra RNA library preparation kit (New England BioLabs) served for NGS library preparation and indexing of the samples, and the NEBNext Library Quant kit for Illumina (New England BioLabs) allowed quantification of the libraries before sequencing on the Illumina MiSeq sequencer (Illumina) using the MiSeq reagent kit (version 3; Illumina). Lazypipe (41) served for *de novo* assembly of SwSCV-1 genome from the samples. In parallel, reference assembly using Bowtie2 (42) run in Unipro UGENE (36) served to obtain consensus sequences for the SwSCV-1 from different cell lines.

To confirm the presence of the mutation in ORF2, we isolated RNA from I/1Ki-mutORF2Δ cells at approximately two months post transfection, and synthesized cDNA using the RevertAid First Strand cDNA Synthesis Kit (ThermoFisher Scientific) according to the manufacturer’s recommendation for random hexamers. The following two primer pairs: FWD-1 5’-CTTTCCGGTACCCCTTGAGT-3’, REV-1 5’ GAACAATGGGTCGAGCTTGG-3’, FWD-2 5’-CTTCCCCAGCTCCTCCATAG-3’, REV-2 5’-AGATCCCTACTCCGTCCAGA-3’ and the Phusion Flash High-Fidelity PCR Master Mix (ThermoFisher Scientific) used according to the manufacturer’s recommendation served to amplify the region containing the introduced mutation. Sanger sequencing (performed by the DNA Sequencing and Genomic Laboratory, Institute of Biotechnology, University of Helsinki) served to confirm the presence of the ORF2 mutation.

For the strand-specific RNA sequencing, we isolated RNA from a 75 cm^2^ cell culture flask of 1.2× SwSCV-1 FWD transfected cells (4 weeks post transfection) using TRIzol reagent (Invitrogen) according to the manufacturer’s recommendation. We split the extracted RNA into two aliquots, which we sent to Azenta Life Sciences for sequencing. Both poly(A) enrichment and rRNA depletion were utilized for RNA selection prior to library preparation. We then aligned to the SwSCV-1 genome and antigenome the reads representing the direction of the original RNA template using Bowtie2 in Ugene.

## RESULTS

### Mutation of the ORF2 does not inhibit initiation of replication

To investigate the potential role of the putative protein encoded by SwSCV-1 ORF2, we established an infectious clone with a single point mutation turning the methionine initiation codon into isoleucine codon while maintaining the amino acid sequence of the DAg. The insert, designed according to the previously described 1.2× genome copy of SwSCV-1 approach (27) is schematically depicted in **Figure 1A**. As described for 1.2× and 2× genome SwSCV-1 infectious clones in earlier studies (17, 27), we generated constructs with the insert in both genomic (mutORF2 SwSCV-1 FWD) and antigenomic (mutORF2 SwSCV-1 REV) orientation. Based on our earlier studies, transfection of the FWD construct to the boa constrictor kidney cell line, I/1Ki, leads to DAg expression only in the case of virus replication, while the REV construct will express the protein also through the CAG promoter of the plasmid (17, 27). To test the ability of the mutORF2 clones to initiate SwSCV-1 replication, we transfected I/1Ki cells with both constructs, fixed or collected cells 1 to 4 days post transfection (dpt) and performed IF staining and WB (**Figure 2A and 2B**). Both approaches displayed increasing SwSCV-1 DAg expression over the course of the four days for both constructs. WB comparison to the 1.2× SwSCV-1 FWD construct showed comparable migration patterns and increasing DAg expression (**Figure 2C**), similarly to our previous observations (27). To further compare the mutORF2 constructs to the wild-type constructs, we transfected I/1Ki cells with all six clones (2×, 1.2× and mutORF2 SwSCV-1 FWD and REV) and collected cell pellets at 1, 2, 3 and 7 dpt. We quantified the amount of SwSCV-1 RNA at each time point by qRT-PCR after DNAse I treatment to remove potential plasmid DNA carryover. **Figure 2D** shows an increasing amount of SwSCV-1 RNA in the transfected cells, indicating replication initiation from all of the constructs at roughly equal efficiency, yielding similar SwSCV-1 RNA levels by day 7.

**Figure 2.**
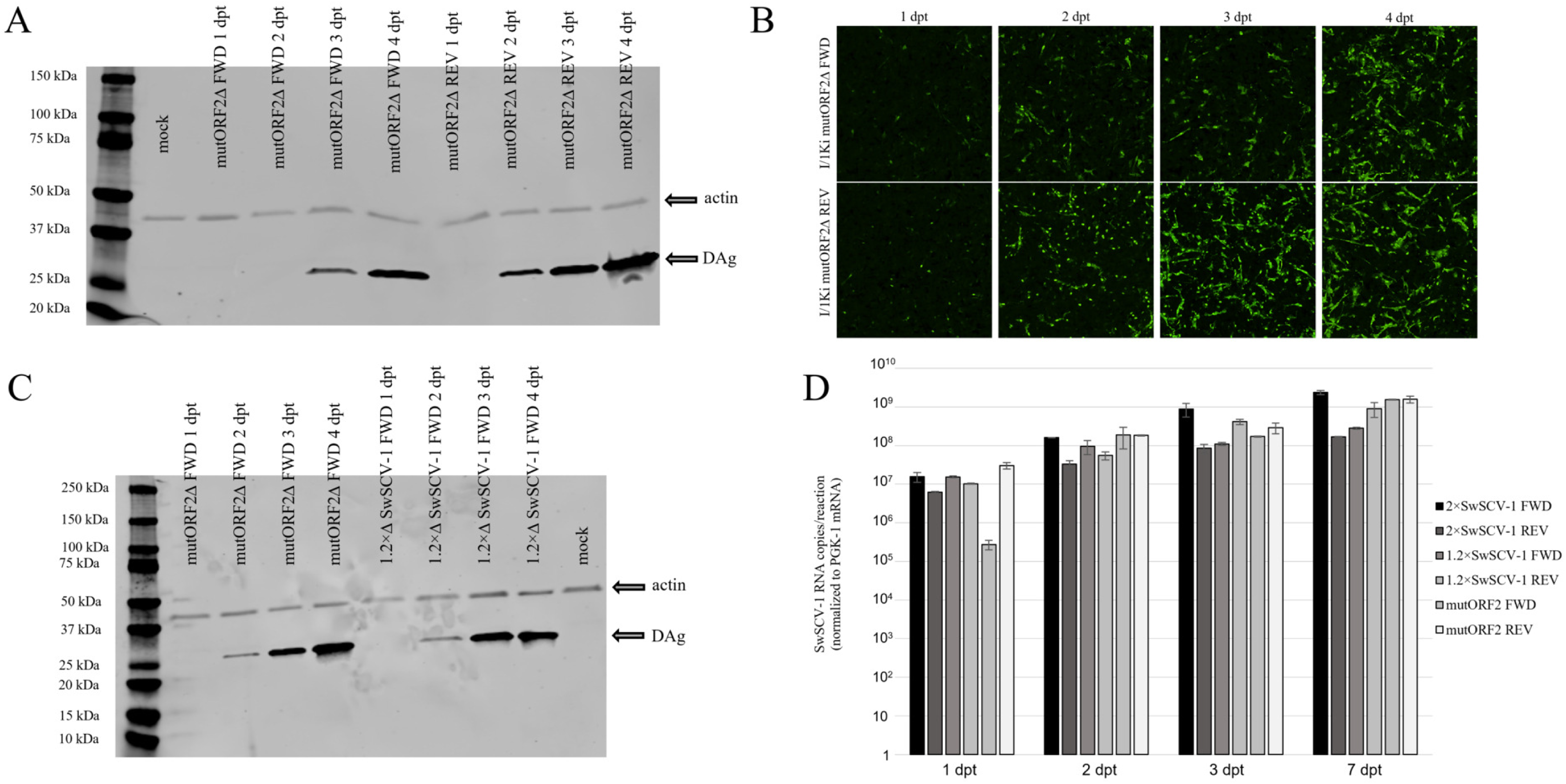
Transfection of mutORF2 SwSCV-1 constructs initiates replication in boa kidney cells. **A)** I/1Ki cell pellets were collected 1 to 4 days post transfection with the mutORF2 SwSCV-1 FWD and REV constructs. The samples were separated on 4–20% Mini-PROTEAN TGX gels, transferred onto nitrocellulose membrane, which were probed with rabbit α-SwSCV-1 DAg antiserum and mouse monoclonal anti-pan actin antibody. The results were recorded using Odyssey Infrared Imaging System. **B)** I/1Ki cells were fixed and stained for the DAg with rabbit α-SwSCV-1 DAg antiserum at 1 to 4 days post transfection with the mutORF2 SwSCV-1 FWD and REV constructs. AlexaFluor 488-labeled donkey anti-rabbit IgG served as the secondary antibody. The images were taken by the Opera Phenix High Content Screening System with 10× objective. **C)** Western blot comparison of DAg expression 1 to 4 days following transfection with the mutORF2 SwSCV-1 FWD and the previously described 1.2× genome SwSCV-1 FWD (27) constructs. The membranes were probed with rabbit α-SwSCV-1 DAg antiserum and mouse monoclonal anti-pan actin antibody and the results were recorded using Odyssey Infrared Imaging System. **D)** I/1Ki cells were transfected with 2×, 1.2× genome and mutORF2 SwSCV-1 FWD and REV constructs, followed by RNA isolation 1-3 and 7 days post transfection. The samples were analyzed by qRT-PCR after DNase I treatment, targeting genomic SwSCV-1 RNA. The results were normalized to PGK-1 mRNA. The y-axis shows SwSCV-1 RNA copy numbers/reaction and the error bars represent standard deviation.

### ORF2 and its putative protein are not essential for persistent infection or infectious particle production

In our previous studies (17, 27), we observed that maintaining the cells after the initial transfection with the 2× SwSCV-1 FWD and 1.2× SwSCV-1 FWD constructs resulted in persistent infection of the cell line. To study if the ORF2 mutation would inhibit the establishment of persistent infection, we transfected I/1Ki cells with the mutORF2 SwSCV-1 FWD construct and passaged the cells to expand the culture, an approach earlier shown to result in persistent infection and loss of the original plasmid (17, 27). IF and WB analyses of the cells revealed continued DAg expression, which based on our earlier studies we interpret to indicate persistent infection, implying that the putative ORF2 protein is not essential for replication. Roughly one year after the initial transfection, we compared the DAg expression and SwSCV-1 RNA levels in the three persistently SwSCV-1 infected cell lines (I/1Ki-2×Δ, I/1Ki-1.2×Δ, and I/1Ki-mutORF2Δ) by IF staining, northern blot, qRT-PCR, and WB (**Figure 3**). IF staining of the I/1Ki-mutORF2Δ cells showed a similar level of infection in each of the cell lines (**Figure 3A**). Investigation of RNA levels and DAg expression by northern blot, qRT-PCR and WB did not reveal overt differences between the three persistently infected cell lines either (**Figure 3B-D**).

**Figure 3.**
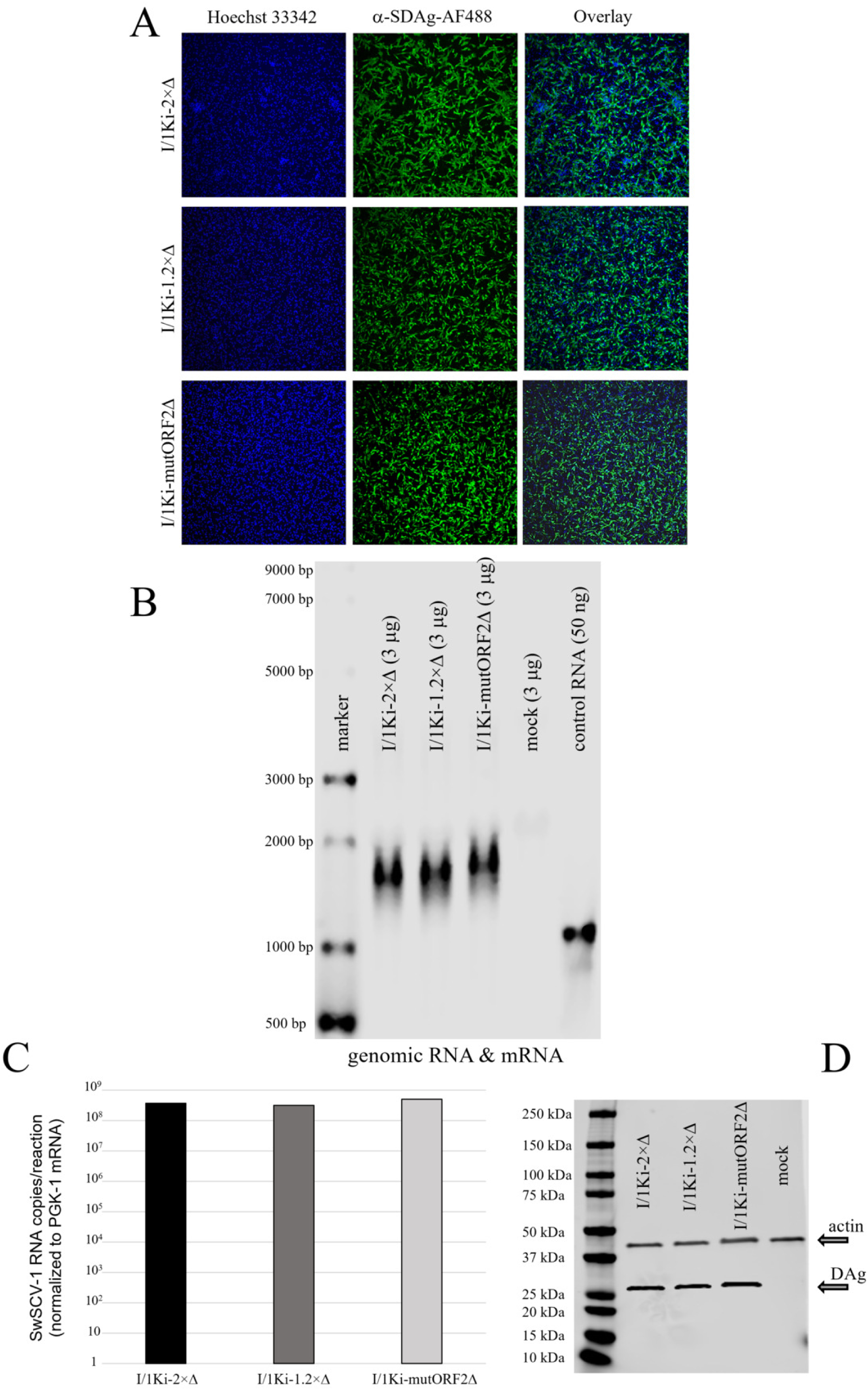
Comparison of the three persistently SwSCV-1 infected cell lines (I/1Ki-2×Δ, I/1Ki-1.2×Δ and I/1Ki-mutORF2Δ). **A)** The persistently infected cell lines were fixed and stained using rabbit α-SwSCV-1 DAg antiserum and Alexa Fluor 488-labeled donkey anti-rabbit secondary antibody, while Hoechst 33342 served for staining the nuclei. The images were taken 3.5 years (I/1Ki-2×Δ), 14 months (I/1Ki-1.2×Δ) and 1.5 years (I/1Ki-mutORF2Δ) post transfection using the Opera Phenix High Content Screening System with 10× objective. **B)** Total RNA was isolated from I/1Ki-2×Δ, I/1Ki-1.2×Δ, I/1Ki-mutORF2Δ and clean I/1Ki cells. An *in vitro* transcribed control RNA (∼850 nucleotides long) served as control. The RNA samples were separated on agarose-formaldehyde gel and transferred onto nylon membrane, which were probed for SwSCV-1 genomic RNA and SwSCV-1 DAg mRNA. The Odyssey Infrared Imaging System served for recording the results. **C)** RNA was isolated from I/1Ki-2×Δ, I/1Ki-1.2×Δ, and I/1Ki-mutORF2Δ cells and analyzed by qRT-PCR targeting SwSCV-1 genomic RNA. The results were normalized to PGK-1 mRNA. The y-axis shows SwSCV-1 RNA copy numbers per reaction. **D)** Samples of I/1Ki-2×Δ, I/1Ki-1.2×Δ, I/1Ki-mutORF2Δ and clean I/1Ki cells were separated on 4–20% Mini-PROTEAN TGX gels, transferred onto nitrocellulose membranes, and probed with rabbit α-SwSCV-1 DAg antiserum and mouse monoclonal anti-pan actin antibody. The results were recorded using Odyssey Infrared Imaging System.

To test if the mutation introduced in the ORF2 would interfere with infectious particle formation, we superinfected all three cell lines three weeks (thereby representing acute SwSCV-1 infection) after the initial transfection (2×, 1.2× and mutORF2 SwSCV-1 FWD) with Haartman Institute snake virus 1 (HISV-1, a hartmanivirus) a potent helper virus of SwSCV-1 (17). Subsequently, we collected SNTs from the superinfected cells up to nine days post infection (dpi) and used them to inoculate naïve I/1Ki cells, which we analyzed by IF staining at 4 dpi for DAg expression. The IF staining (**Figure 4A**) of the cells inoculated with the SNTs collected from the superinfected cells at 3, 6, and 9 dpi showed similar staining patterns and similar levels of infectious particles. Quantification of the infectious SwSCV-1 particles formed after superinfection of the three cell lines confirmed the numbers of infectious particles secreted from all cell lines to be in the same range (**Figure 4B**).

**Figure 4.**
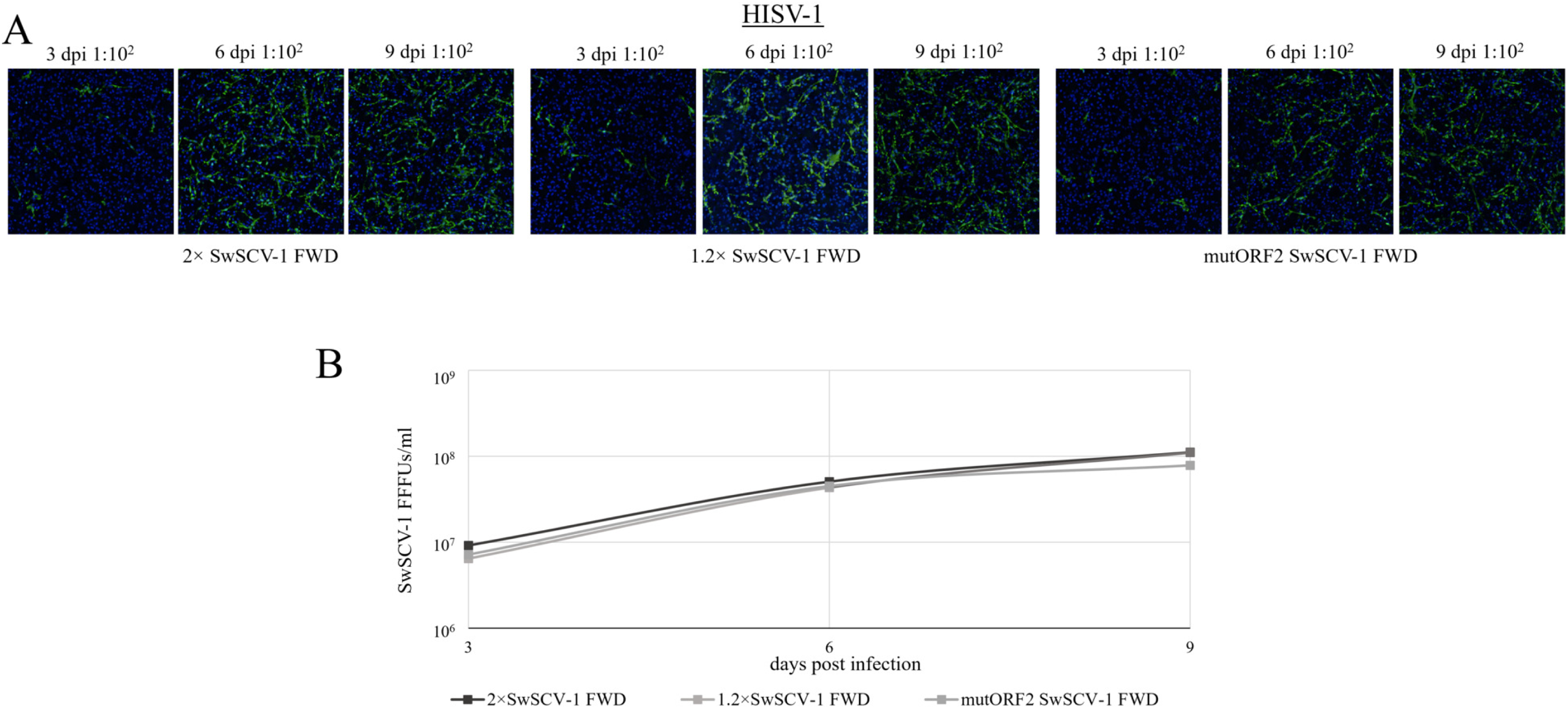
Comparison of the infectious particle production of the three freshly SwSCV-1 transfected (2×, 1.2× and mutORF2 SwSCV-1 FWD) cell lines. **A)** The three freshly transfected I/1Ki cell lines were superinfected with HISV-1, supernatants were collected 3, 6 and 9 days post infection and subsequently used for titration on naïve I/1Ki cells. Four days post inoculation; we fixed and stained the cells using rabbit α-SwSCV-1 DAg antiserum and Alexa Fluor 488-labeled donkey anti-rabbit secondary antibody, while Hoechst 33342 served for staining the nuclei. The images were taken by the Opera Phenix High Content Screening System with 10× objective. **B)** The number of infected cells was counted using the Opera Phenix High Content Screening System, followed by quantification of infectious particles per milliliter of growth medium in terms of fluorescent focus-forming units (FFFUs—displayed on y-axis).

### Passaging of SwSCV-1 infected cells leads to re-establishment of the ORF2

To study if persistent infection would induce mutations in the SwSCV-1 genome, we performed RNA sequencing of the three persistently SwSCV-1 infected cell lines (I/1Ki-2×Δ, I/1Ki-1.2×Δ, and I/1Ki-mutORF2Δ) analyzed >1 year after initiation of infection through transfection. The consensus sequences generated using Bowtie2 (42) from the RNA sequencing reads showed the presence of wild type SwSCV-1 genome in the I/1Ki-1.2×Δ cells. However, the consensus sequence generated from the I/1Ki-2×Δ cells revealed a T to C point mutation at position 238 (GenBank accession no. MH988742.1) (TGGACTCCGGCGTTACTCGCATGGG T→C) in the SwSCV-1 genome. The same position showed high T to C variation also in the case of I/1Ki-1.2×Δ and I/1Ki-mutORF2Δ cell lines, and interestingly the mutation would introduce a methionine codon in the antigenomic sense (CGTAT→CG<u>C</u>AT). The resulting ORF would only be ∼15 amino acids long, thereby likely not representing a true protein. Curiously, the reference assembly of the SwSCV-1 genome from the I/1Ki-mutORF2Δ cell line showed that the introduced mutation had fully reverted during persistence (>1 year of infection), re-introducing the methionine initiation codon of the ORF2. When we analyzed the transfected cells by RT-PCR and Sanger sequencing at approximately two months post initial transfection, we did not observe the reversion, likely due to its low abundance at that point.

### The putative protein product of ORF2 is cytoplasmic and SwSCV-1 infected snake does not have a detectable ORF2 antibody response

After observing reversion of the mutation in the ORF2 during persistent infection, we decided to investigate if we could detect the putative ORF2 protein product in SwSCV-1 infected I/1Ki cells. We attempted generating rabbit antiserum against his-tagged ORF2 protein product expressed in bacteria, as well as two selected peptides of the putative protein. Both approaches failed to produce an antiserum suitable for WB detection of the putative ORF2 encoded protein expressed in I/1Ki cells from constructs with and without C-terminal HA-tag. Utilizing anti-HA antibody, we observed boiling and/or addition of reducing agent (DTT) to render the ORF2 protein undetectable in WB. After identifying a suitable sample pre-treatment, WB analysis of I/1Ki cell pellets collected three days post ORF2-HA and ORF2 expression construct transfection, we found the putative protein to migrate at a rate approximately similar to the 20 kDa molecular weight marker (**Figure 5A**). We hypothesized that if the protein is expressed at high levels during infection, an infected animal would produce antibodies against the ORF2 protein product. To test this hypothesis, we prepared a WB membrane of I/1Ki cells transfected with wild-type and mutORF2 SwSCV-1 (1.2× SwSCV-1 FWD and mutORF2 SwSCV-1 FWD) infectious clones, persistently SwSCV-1 infected (I/1Ki-1.2×Δ) I/1Ki cells, and I/1Ki cells transfected with the ORF2 expression constructs (with or without HA-tag), and probed the membrane with the serum of the original SwSCV-1 infected boa constrictor (16). The results showed that the snake serum indeed reacted with the DAg in all SwSCV-1 transfected/infected samples. However, it did not react with the ORF2 produced in I/1Ki through recombinant expression (**Figure 5A**). The snake serum did not produce a similar sized band in any of the transfected/infected cell lysates either, suggesting that ORF2 would not be expressed during SwSCV-1 infection or that the putative protein would be poorly immunogenic and/or expressed at very low level.

**Figure 5.**
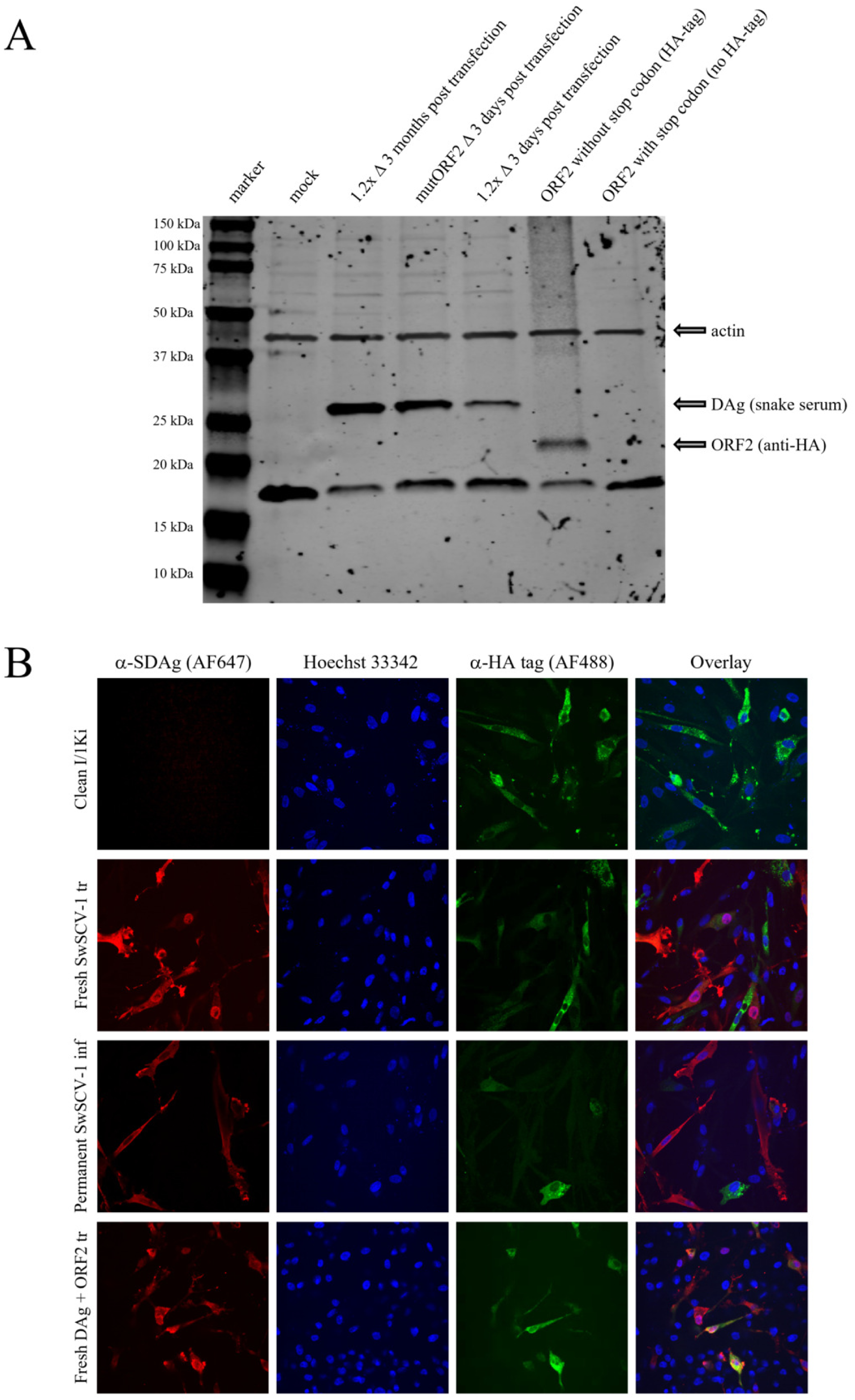
Comparing ORF2 expression patterns in freshly or persistently SwSCV-1 infected cells versus in cells transfected with an ORF2 expression construct by western blot and immunofluorescence staining. **A)** Samples of persistently SwSCV-1 infected, freshly SwSCV-1 transfected, ORF2 (with or without an HA-tag) expression construct transfected, and clean I/1Ki cells were separated on 4–20% Mini-PROTEAN TGX gels, transferred onto nitrocellulose membranes. The membrane was probed with the serum of a SwSCV-1 infected snake at 1:250 dilution (16), anti-HA antibody at 1:1000 dilution, and mouse monoclonal anti-pan actin antibody. The results were recorded using Odyssey Infrared Imaging System. **B)** Recombinant expression of the ORF2-HA protein in clean I/1Ki cells (top row), freshly SwSCV-1 transfected cells (second row), persistently SwSCV-1 infected cells (third row), and snake DAg transfected cells (bottom row). The ORF2 protein shows cytoplasmic subcellular localization with a granular staining pattern and does not co-localize with the DAg. The cells were fixed and stained using rabbit α-SwSCV-1 DAg antiserum with Alexa Fluor 647-labeled donkey anti-rabbit secondary antibody, mouse α-HA primary antibody with Alexa Fluor 488-labeled donkey anti-mouse secondary antibody, while Hoechst 33342 served for staining the nuclei. The images were taken using the Opera Phenix High Content Screening System with 63× objective.

To further study the putative protein product of ORF2, we transfected I/1Ki cells with expression constructs for HA-tag bearing ORF2 and used IF staining to study the subcellular localization of the protein in non-infected, SwSCV-1 infected, or SwSCV-1 DAg transfected I/1Ki cells. The IF staining showed a fairly intense granular or grainy staining pattern in the cytoplasm for the putative ORF2 protein in the uninfected cells (**Figure 5B**). However, in cells infected with SwSCV-1 for <4 months (1.2× SwSCV-1 FWD), persistently (>4 months) SwSCV-1 infected (I/1Ki-1.2×Δ) cells, or cells co-transfected with SwSCV-1 DAg, the ORF2 protein staining was considerably weaker. These observations could imply competition between the plasmid and the virus from the cellular RNA polymerase II’s resources or ORF2-DAg interaction mediated relocation, as ORF2 and DAg did not appear to colocalize (**Figure 5B**).

### Directional RNAseq suggests that ORF2 is not transcribed to mRNA

To investigate the transcriptional patterns of SwSCV-1 in more detail, we performed strand-specific RNAseq on I/1Ki cells transfected with the 1.2× SwSCV-1 FWD infectious clone. The method preserves the polarity of the transcripts and allows to determine the orientation of the reads and to identify the mRNA species within the cells. We utilized two approaches for selecting the RNA species for library preparation, poly(A) enrichment and ribosomal RNA (rRNA) depletion. Poly(A) enrichment focuses on mRNA species with polyadenylation, whereas rRNA depletion offers a more comprehensive representation of cellular RNA, including non-coding transcripts. We aligned the reads representing the original RNA template orientation to the 1.2× SwSCV-1 genome using Bowtie2 to allow assessment of the reads spanning the circular junctions of the genome. The genome is the most abundant viral RNA species during HDV replication (4), and indeed the mapping showed a strong bias towards the genomic strand in the case of reads obtained following rRNA depletion (**Figure 6A**). The reads obtained from poly(A) enriched RNA showed the presence of DAg mRNA (**Figure 6B**) with a small bias towards the 5’ end of the mRNA. We did not locate a classical AAUAAA polyadenylation signal following the ORF2, however, there are both AAGAAA and GATAAA signals, these can theoretically be utilized for polyadenylation (43). Neither the rRNA depletion nor poly(A) enrichment of the RNA prior to library preparation produced an obvious increase in reads covering the ORF2 (**Figure 6**). In addition to DAg mRNA, we observed a high amount of reads mapping to the opposing side of the rod (**Figure 6B**), perhaps mediated through co-purification of the opposing RNA strand (reflecting the self-complementary structure of kolmivirus) rather than direct polyadenylation mediated enrichment of this region.

**Figure 6.**
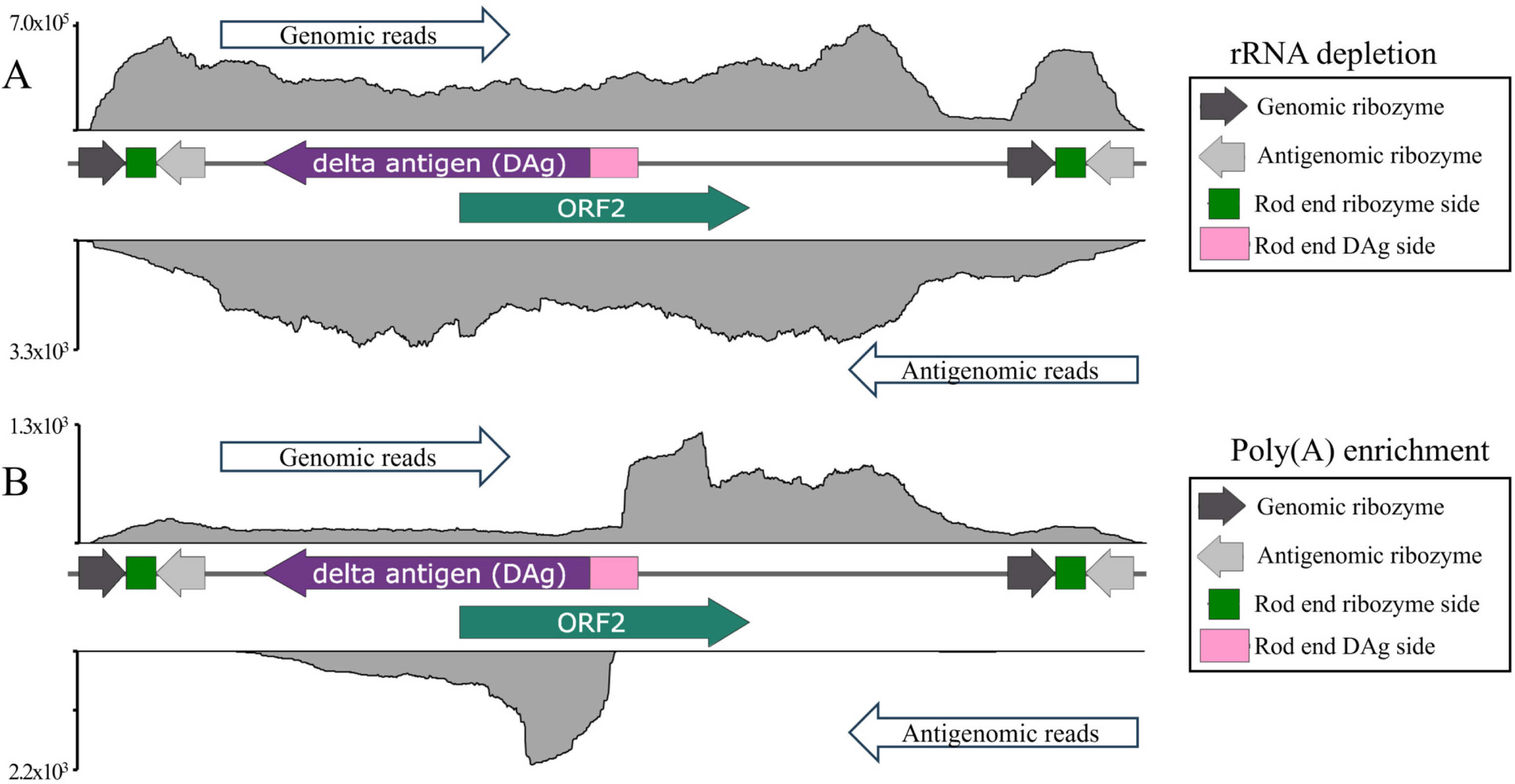
Directional RNAseq coverage maps of I/1Ki cells four weeks post transfection with the 1.2× SwSCV-1 FWD infectious clone. Libraries were prepared using either rRNA depletion or poly(A) enrichment. The obtained reads were aligned to the 1.2× SwSCV-1 genome using Bowtie2. The y-axis shows the number of genomic and antigenomic mapped reads. **A)** rRNA depleted libraries show a strong bias towards the genomic strand, consistent with the genome being the most abundant RNA species during kolmiovirid replication. **B)** Poly(A) enriched libraries show reads corresponding to the delta antigen (DAg) mRNA, while coverage over the open reading frame 2 (ORF2) indicates low or absent mRNA transcription of the region.

### SwSCV-1 does not express the large form of the delta antigen

To further characterize the SwSCV-1 genome, we investigated whether its DAg has two forms. In the case of HDV, the two forms of DAg, S- and L-DAg, mediate different functions (12). The HDV S-DAg prominently localizes to the nucleus and is critical for replication (44) whereas the L-DAg associates with particle formation (45). In our earlier studies, when investigating infected tissues and cultured cells, we observed two (or more) bands in WBs when staining for SwSCV-1 DAg (16, 17), and speculated that the two bands could correspond to the S- and L-DAg, post-translationally modified forms of the DAg, or proteolytic degradation of the DAg. We generated three constructs for recombinant expression of the SwSCV-1 DAg, one with the unedited SwSCV-1 DAg ORF, the second with S-DAg ORF only, and the third with DAg ORF in which the stop codon is replaced by tryptophan codon forcing L-DAg translation (**Figure 1C**). We transfected I/1Ki cells with the constructs and analyzed the protein expression by IF staining. This demonstrated similar if not identical staining patterns for the three DAg expression constructs (**Figure 7A**). We then compared the electrophoretic mobility of the DAg produced via plasmid expression to DAg produced by the persistently SwSCV-1 infected (**Figure 7B**) and freshly SwSCV-1 transfected cell lines (**Figure 8A**). The WB showed that both the freshly infected cells and the persistently infected cell lines only expressed the S-DAg form. This suggests that SwSCV-1 only expresses and requires a single DAg form for replication and infectious particle formation. To study whether the L-DAg expression occurs in cell lines of different tissue or host origin, we used the persistently SwSCV-1 infected snake cell lines described earlier (V/5Lu-Δ, V/1Ki-Δ, V/2Hz-Δ, and V/5Liv-Δ) (17) and a recently generated one (V/4Br-Δ), as well as two human cell lines (Huh7 and HEK293T) and compared their DAg expression by WB. The results show that regardless of the tissue or host origin of the cell line, there is only a single band, corresponding to the S-DAg (**Figure 8B-C**).

**Figure 7.**
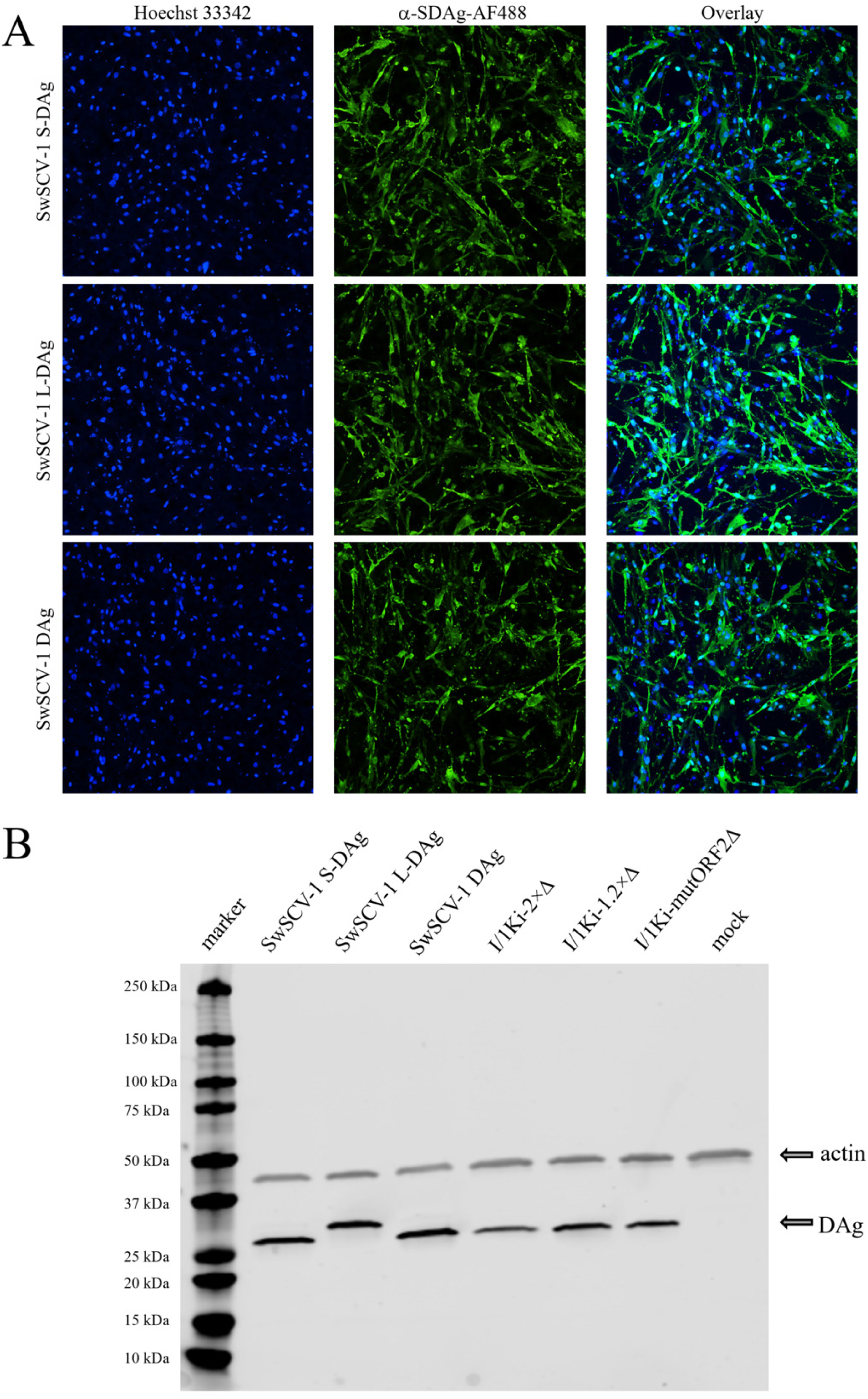
Comparison of protein expression after transfection with expression constructs versus infectious clones. **A)** I/1Ki cells were transfected with plasmids bearing the small SwSCV-1 DAg (top panels), large SwSCV-1 DAg (second row panels), the unedited SwSCV-1 DAg (third row panels). The cells were fixed and stained 3 days post transfection using rabbit α-SwSCV-1 DAg antiserum and Alexa Fluor 488-labeled donkey anti-rabbit secondary antibody, while Hoechst 33342 served for staining the nuclei. The images were taken using the Opera Phenix High Content Screening System with 20× objective. **B)** I/1Ki cells 3 days post transfection with SwSCV-1 S-DAg, L-DAg and unedited DAg bearing construct and I/1Ki-2×Δ, I/1Ki-1.2×Δ and I/1Ki-mutORF2Δ cells were separated on 4–20% Mini-PROTEAN TGX gels, transferred onto nitrocellulose membranes, followed by probing with rabbit α-SwSCV-1 DAg antiserum and mouse monoclonal anti-pan actin antibody. The results were recorded using Odyssey Infrared Imaging System.

**Figure 8.**
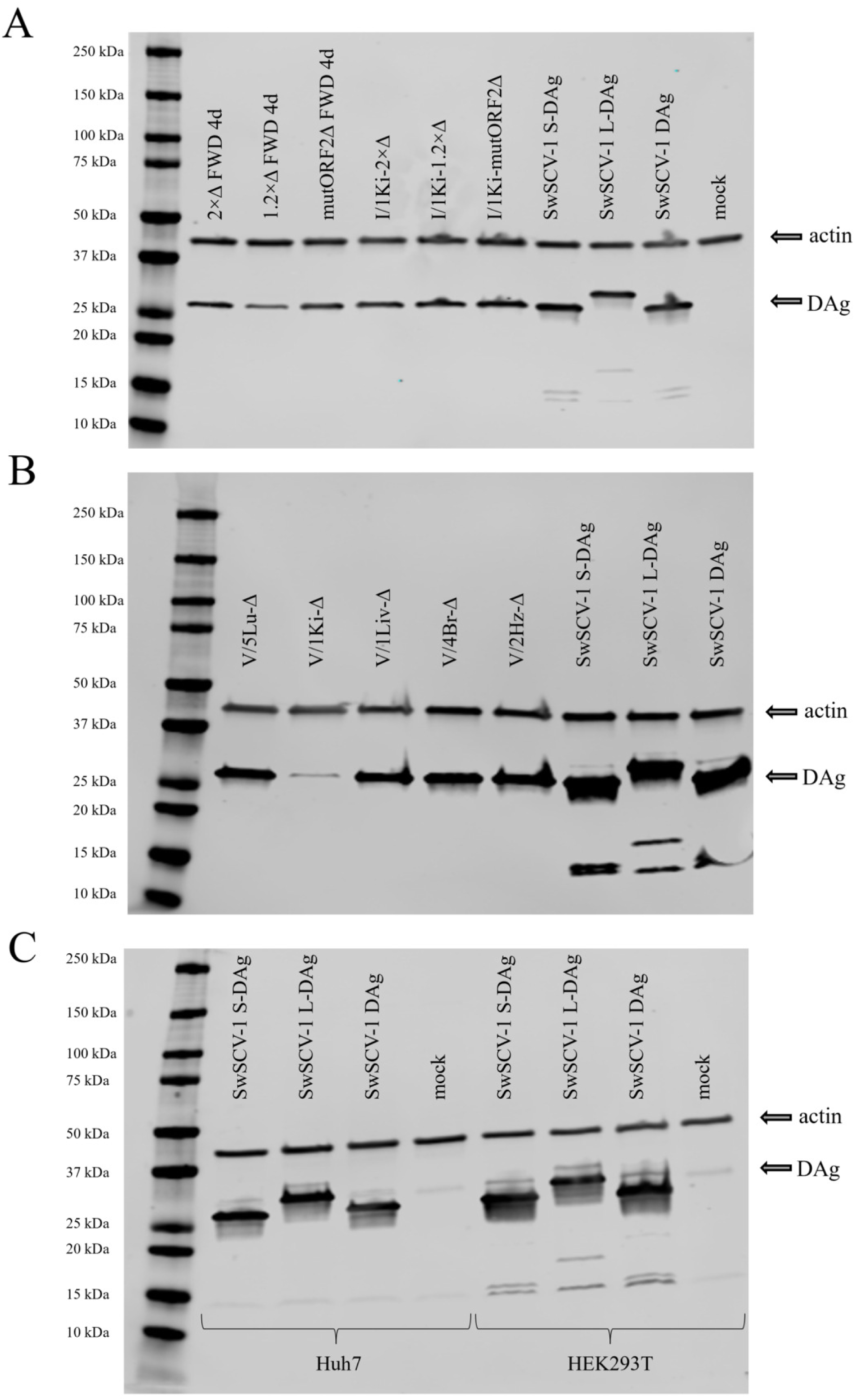
SwSCV-1 does not express large form of the delta antigen. **A)** Western blot comparison of freshly SwSCV-1 transfected cells, persistently SwSCv-1 infected cells, and I/1Ki cells transfected with the SwSCV-1 S-DAg, L-DAg and unedited DAg expression constructs. **B)** Western blot comparison of persistently SwSCV-1 infected cell lines originating from various tissues of boa constrictor snakes and cells transfected with the SwSCV-1 S-DAg, L-DAg and DAg expression constructs **C)** Western blot comparison of human hepatocellular carcinoma (Huh7) and human embryonic kidney (HEK293T) cells transfected with the SwSCV-1 S-DAg, L-DAg and DAg expression constructs. Cell pellets were collected 3 days post transfection. (**A**, **B** and **C**) The proteins were separated on 4–20% Mini-PROTEAN TGX gels, followed by transfer onto nitrocellulose membranes. The membranes were were probed with rabbit α-SwSCV-1 DAg antiserum and mouse monoclonal anti-pan actin antibody.

## DISCUSSION

Owing to almost half a century of research since the discovery of HDV (1), a wealth of knowledge on the unique features of HDV is available. However, the newly-identified distinct HDV-like agents from a wide range of taxa (15, 16, 18–21), classified in different species within the family *Kolmioviridae* of the realm *<u>Ribozyviria</u>* (23, 24), have brought new dimensions to the research around these enigmatic agents. So far, however, reports have characterized only some of the novel agents further than genome description, calling for studies comparing them against the best-known member, HDV. After identifying SwSCV-1, we have been developing molecular biology tools that allow us to study SwSCV-1 more extensively. So far, we have shown its ability to initiate replication *in vitro* following the transfection of a dimer or 1.2× genome constructs (17, 27), in a similar manner to HDV (46–48). In the present study, we focused on the ORFs present in the SwSCV-1 genome. For HDV, the early studies showed that during replication only the DAg ORF appears to be translated (5). A recent interesting study indicated that an additional protein would be translated from the *Serinus canaria*-associated deltavirus genome (26), further emphasizing the importance of investigating the novel kolmiovirids in more depth. The genome of SwSCV-1 harbors two partially overlapping >500 base ORFs, the DAg ORF in the antigenomic orientation and the ORF2 in the genomic orientation (**Figure 1A**). The putative protein encoded by ORF2 does not show substantial similarity to any known protein. Since the sequence of the putative *Serinus canaria*–associated deltavirus ORF2 is not available, we were unable to perform a comparison. Therefore, we investigated whether disruption of ORF2 affects the replication capacity of SwSCV-1. In addition, we examined whether the SwSCV-1 DAg is expressed in two distinct forms, as reported for HDV (7).

To explore the possible roles of the putative ORF2 protein, we generated a construct in which an isoleucine codon replaces the “wild-type” methionine initiation codon of the SwSCV-1 ORF2. By comparing the “mutORF2” construct to wild-type SwSCV-1 genome-bearing constructs, we showed that eliminating the possibility of protein translation does not inhibit initiation of replication or the ability to establish persistent infection (**Figures 2 and 3**). Our results also showed that the introduced mutation did not markedly affect the ability of infectious SwSCV-1 particle formation upon superinfection with a potent helper virus (**Figure 4**).

RNA sequencing of the persistently SwSCV-1 infected cells did not reveal significant changes in the genome of the virus. We identified a single nucleotide mutation in the consensus sequence of SwSCV-1 obtained from cells transfected with the with 2× SwSCV-1 FWD construct approximately three and a half years earlier. The other persistently infected cell lines showed variation at the same sequence position, but the altered nucleotide had not yet become dominant in those cell lines. However, both the wild-type and the mutated sequence appeared abundant in the genome of SwSCV-1 obtained from all of the persistently infected cell lines. It seems unlikely that the single nucleotide mutation outside the DAg ORF and the ORF2 would dramatically affect the viral life cycle. On the other hand, the point mutation could alter the pairing between the opposite sides of the circular single-stranded RNA genome, thereby affecting the transcription. The sequencing further revealed that the mutation introduced into the ORF2 initiation codon had reverted in the persistently infected cells. The reversion was not present in the freshly transfected cells analyzed two months after transfection by Sanger sequencing, suggesting the reversion to be beneficial for the virus. We speculate that the introduced nucleotide change “de-optimized” the respective amino acid codon in the DAg ORF. The nucleotide change from AG<u>C</u>AT to AG<u>T</u>AT changes a serine codon of the DAg ORF from AGC to AGT that in the case of *Python molurus bivitattus* (codon usage table for boa constrictor not available) is approximately equally abundant (Codon Usage Database, https://www.kazusa.or.jp/codon/).

By transfecting ORF2 expressing constructs into boa kidney cells, we were able to confirm that a stable protein can be translated from the ORF. Although we were unsuccessful in our trials to generate an ORF2 specific antibody, the addition of an HA-tag into the protein expressing constructs allowed subsequent detection of ORF2 both in IF staining and WB. Co-expression of the putative protein and DAg did not alter the staining pattern of either protein nor did they co-localize, suggesting that the proteins would be unlikely to interact. The serum of the snake in which we originally identified SwSCV-1 (16) failed to detect the putative ORF2-encoded protein, suggesting that the protein is not expressed during the viral life cycle or that no antibodies are produced against it due to, e.g., low immunogenicity, very low expression levels, or a complete lack of its expression.

Directional RNA sequencing of cells transfected with 1.2× SwSCV-1 FWD and analyzed at four weeks post transfection indicated a transcript from the DAg ORF as expected. During poly(A) selection in library preparation mRNAs are captured by their 3’ end poly(A) tail, causing a possible bias in the amount of reads towards the 3’ end of the transcript, which we could observe in the case of the DAg. In contrast, the sequencing data could not convincingly demonstrate the presence of ORF2 mRNA (**Figure 6B**), however, it does not completely rule out its possible presence. The kolmiovirus genomes form rod-like structures through extensive internal base pairing of the circular single-stranded genome, and the ORF2 spans the “rod-end” of the genome. Considering the fact that the “rod-end” of the genome end acts as the transcriptional promoter of HDV (49, 50), it would seem unlikely that an actively employed ORF would span this region. The ORF2 lacks a downstream conventional poly(A) signal, however, there are two non-conventional poly(A) signals present downstream of ORF2 which could imply that the protein product of this ORF can be translated in some cells or tissues. Interestingly, poly(A) selection revealed reads mapping to the SwSCV-1 rod on the opposing side of the DAg ORF (**Figure 6B**). This could indicate that similarly to HDV (50), the rod end functions as a bidirectional promoter. These observations would merit further research.

In the case of HDV, the S- and L-DAg carry different roles during the viral life cycle (7, 8, 12). In our earlier studies, using WB, we have observed that SwSCV-1 DAg migrates mainly as a single species, however, we were not able to conclude whether the majority of the protein would be in the S- or L-DAg form. Herein, using recombinant proteins, we could conclude that the SwSCV-1 infected I/1Ki cells mostly, if not solely, produce S-DAg (**Figure 7B** and **Figure 8A**). This would be in line with the findings that some other non-HDV kolmioviruses only express a single form of the DAg (21, 51, 52). In addition, Wille and co-authors did not, based on sequence analysis, find a putative ADAR editing site in the DAg of dabbling duck virus 1 (15). Iwamoto and colleagues found no possible editing site in the sequences of *Taeniopygia guttata* and *Marmota monax* deltaviruses by looking at nucleotide variation, while they identified a potential RNA-editing site in the sequence of *Odocoileus virginianus* deltavirus, however, the elongation would be only with two amino acids (19). Tome’s spiny rat virus likewise contains an amber stop codon, suggesting the potential for ADAR-mediated editing (52). However, neither knocking out nor overexpressing of the ADAR altered the expression pattern of the rodent DAg as assessed by WB analysis (52). Because we had in our earlier studies observed double or multiple bands for DAg in WB of some samples (16, 17, 27), we further studied the possibility that the ADAR editing resulting in L-DAg translation would occur in cell lines originating from different tissues. WB of different snake cell lines infected with SwSCV-1 and of human cell lines transfected with the SwSCV-1 DAg expression constructs all showed only the presence of S-DAg (**Figure 8B-C**), suggesting that the expression levels or lack of ADAR expression in different cell types would not affect the S- versus L-DAg in the case of SwSCV-1. We currently lack the tools to confirm ADAR expression or to assess its expression levels in boa constrictor–derived cell lines. Moreover, RNA sequencing data from the I/1Ki cells indicate that only a negligible number of reads showed variation at the S-DAg stop codon. The findings would indicate that ADAR editing does not occur in our boa constrictor cell lines under the conditions tested. Furthermore, they may imply that ADAR editing responsible for the elongation of DAg would specifically apply to HDV, and it could relate to co-evolution with HBV in humans.

Although absence of evidence is not evidence of absence, we think that our results provide strong evidence that ORF2 is not translated during infection. Specifically, the experiments with the mutORF2 construct showed that removal of the methionine codon does not markedly affect replication, establishment of persistent infection, or particle formation. However, the methionine codon reappeared during persistent infection, suggesting codon usage bias in SwSCV-1. The absence of antibodies against the putative protein in SwSCV-1 infected animals and the data from the strand-specific RNA sequencing are additional confirmation that no protein is being translated from the ORF2. Even though SwSCV-1 seems to express a single protein, similarly to HDV, the translation of additional ORFs in the *Kolmioviridae* family warrants further studies. By reporting our findings, we hope to highlight the importance of studying the potential of additional ORFs in kolmiovirids, which seems to be a surprisingly overlooked topic in the currently existing literature.

## ACKNOWLEDGEMENTS

The authors wish to acknowledge Antti Hassinen of the FIMM (Institute for Molecular Medicine Finland) High Content Imaging and Analysis (FIMM-HCA) for expert help in imaging and quantification of the IF staining for titration. The study was funded by the Sigrid Jusélius foundation (to JH) and Jane and Aatos Erkko Foundation (to JH), the funding bodies had no role in the study design, interpretation of the results or preparation of the manuscript.

